# Nucleosome wrapping state encodes principles of 3D genome organization

**DOI:** 10.1101/2023.12.20.572694

**Authors:** Zengqi Wen, Xinqian Yu, Ruixin Fang, Haizhen Long

## Abstract

Nucleosome is the basic structural unit of the genome. During processes like DNA replication and gene transcription, the conformation of nucleosomes undergoes dynamic changes, including DNA unwrapping and rewrapping, as well as histone disassembly and assembly. However, the wrapping characteristics of nucleosomes across the entire genome, including region-specificity and their correlation with higher-order chromatin organization, remain to be studied. In this study, we investigated the wrapping length of DNA on nucleosomes across the whole genome using wrapping-seq. We discovered that the chromatin of mouse ES cells forms <u>N</u>ucleosome Wrapping <u>D</u>omains (NRDs), which is conserved in yeast and fly. We found that the degree of nucleosome wrapping decreases after DNA replication and is promoted by transcription. Furthermore, we observed that nucleosome wrapping states delineate 3D genome organization and DNA replication timing. In conclusion, we have revealed a novel principle of higher-order chromatin organization.

## Introduction

In eukaryotic organisms, genomic DNA wraps with histones to form chromatin. The nucleosome is the fundamental structural unit of chromatin. The first high-resolution (2.8-Å resolution) crystal structure of the nucleosome was determined by Luger *et al.* in 1997 ^1^. The structure revealed that a nucleosome core particle contains 147 base pair of DNA wrapping about 1.65 turns around the surface of histone octamer. Multiple interactions are formed within the histone octamer and between the histone octamer and DNA, thus forming a stable complex. Nucleosome is assembled in a stepwise fashion ^2^. An H3-H4 tetramer initially associates with DNA to form a tetrasome, which can then be followed by the addition of one copy of H2A-H2B to form a hexasome. Alternatively, two copies of H2A-H2B can be consecutively added to create a complete nucleosome. Thus, intermediate states, including DNA unwrapping and octamer dissociation, among others may exist during DNA replication when nucleosomes are disassembled ahead of replication fork and re-assembled behind the replication fork ^3^.

As a large complex, nucleosome is dynamic in nature. For instance, DNA located on nucleosomes can rapidly transition between an “unwrapping” and “rewrapping” states, a phenomenon referred to as “DNA breathing”. Utilizing fluorescence resonance energy transfer (FRET), it was demonstrated that DNA on nucleosomes spends approximately 2-10% of its time in the unwrapped state ^4,5^. Significantly, this characteristic is also observed in nucleosomes within nucleosome arrays ^6^. Moreover, the dynamics of nucleosomes within the cell are regulated during various biological processes. Chromatin remodeling complexes are able to regulate the structure of nucleosomes using the energy generated by ATP hydrolysis ^7^. For instance, the nucleosome remodeling complex INO80, can disrupt interactions between H2A and DNA, leading to the opening of approximately 15 base pairs of DNA on the nucleosome ^8,9^. Histone chaperone complexes can bind to histones and regulate the assembly or disassembly of nucleosomes. For instance, the SPT16 subunit of the FACT complex can displace H2A-H2B dimers^10,11^, thus facilitating the formation of an open nucleosome structure.

In the context of gene transcription, RNA polymerase can induce the opening of DNA on the nucleosome, leading to the formation of unwrapped nucleosomes. This process produces intermediate states such as unwrapping DNA of 20 base pairs, 50 base pairs, 60 base pairs, and so forth ^11,12^.

While various nucleosome wrapping states have been observed under *in vivo* conditions, the genuine nucleosome wrapping states within cells remain significantly less characterized. By employing ChIP-exo, Rhee et al. found that histones are asymmetrically depleted on +1 nucleosomes in yeast, leading to the formation of subnucleosomes ^13^. Another widely used approach for mapping nucleosome states is MNase-seq, which directly measures the length of DNA fragments protected by nucleosomes. A recent study from Henikoff’s lab suggested that the protected short DNA fragments associated with nucleosomes can be indicative of subnucleosomal states ^14^. In our previous study, we employed MNase-X-ChIP-seq to map the nucleosome wrapping states in mouse ES cells ^15^. We found that H2A.Z nucleosomes are more unwrapped than canonical nucleosomes, and the wrapping states of H2A.Z nucleosomes is correlated with gene transcription activity ^15^.

As those studies of mapping nucleosome states within cells have primarily focused on analyzing the wrapping states of the +1 nucleosomes, we aim to address the characteristics of nucleosome wrapping states across the entire genome in this study. To this end, we have adapted from our previous MNase-X-ChIP-seq protocol ^15^ and utilized the lengths of DNA fragments generated by MNase enzyme cleavage to calculate quantitative indices of nucleosome wrapping, namely <u>N</u>ucleosome Wrapping <u>S</u>core (NRS) and <u>N</u>ucleosome Wrapping Index (NRI). These two parameters were used to characterize the average wrapping states of nucleosomes within certain genomic regions. To distinguish this experimental and analytical workflow from classic MNase-seq and emphasize the analysis of nucleosome wrapping states, we propose referring to the pipeline as “wrapping-seq“.

Importantly, we revealed that the genomic chromatin of mouse ES cells forms <u>N</u>ucleosome Wrapping <u>D</u>omains (NRDs), including tightly wrapped NRDs (TiNRDs) and loosely wrapped NRDs (LoNRDs). Formation of NRDs is conserved in yeast and fly genomes. TiNRDs and LoNRDs precisely coincide with Hi-C A compartment domains and B compartment domains, respectively. Interestingly, our data suggests that nucleosomes in euchromatin chromatin (i.e. A compartments) exhibit tighter wrapping compared to nucleosomes in heterochromatin. Furthermore, we showed that transcription enhances nucleosome wrapping in genic regions. Thus, this study revealed a novel level of higher-order chromatin organization.

## Results

### Nucleosome wrapping index is robust for nucleosome wrapping state characterization

To study the wrapping states of nucleosomes across genome-wide, we performed wrapping-seq for histone H3 and sequenced the library deeply. In total, 210 M high quality paired-end reads were generated after filtering. Then we use fragments within 50-250 bp to analyze genome-wide nucleosome wrapping states (Fig. S1a). As shown in Fig. 1a, within a genomic window (e.g., 10 kb), when a fragment length break point (X bp) is chosen, <u>N</u>ucleosome Wrapping <u>S</u>core (NRS) is computed as the relative deviation between the number (a) of fragments within X-250 bp and the number (b) of fragments within 50-X bp, expressed as NRS(X) = (a-b)/(a+b). When the break point increased from 80 bp to 160 bp with 10 bp step, the genome-wide NRS generally shifted from 1 to -1 (Fig. 1b, Fig. S1b). However, when the raw NRS of each break point was transformed as Z-score to derive <u>N</u>ucleosome Wrapping Index (NRI), the genome-wide NRIs show highly similar pattern (Fig. 1c, Fig. S1c) and high correlation (Fig. S1d) among different break points. Thus, Z-score transformation eliminates the bias of nucleosome wrapping degree introduced by selecting the break point arbitrarily, and we will use NRI(140) mostly in the following text unless indicated. The value of NRI along the genome reflects relative wrapping state of DNA on the nucleosome. In general, larger NRI value indicates DNA wraps tighter on nucleosomes, thus conferring more protection of MNase digestion; smaller NRI value indicates DNA wraps looser on nucleosomes, thus conferring less protection of MNase digestion. Then we computed NRS(140) and NRI(140) of H3 with multiple bin length. While NRS(140)s show variation with different bin length (Fig. S1e), the pattern of NRI(140)s are rather constant, and highly correlated at genome-wide (Fig. S1e-f).

**Figure 1.**
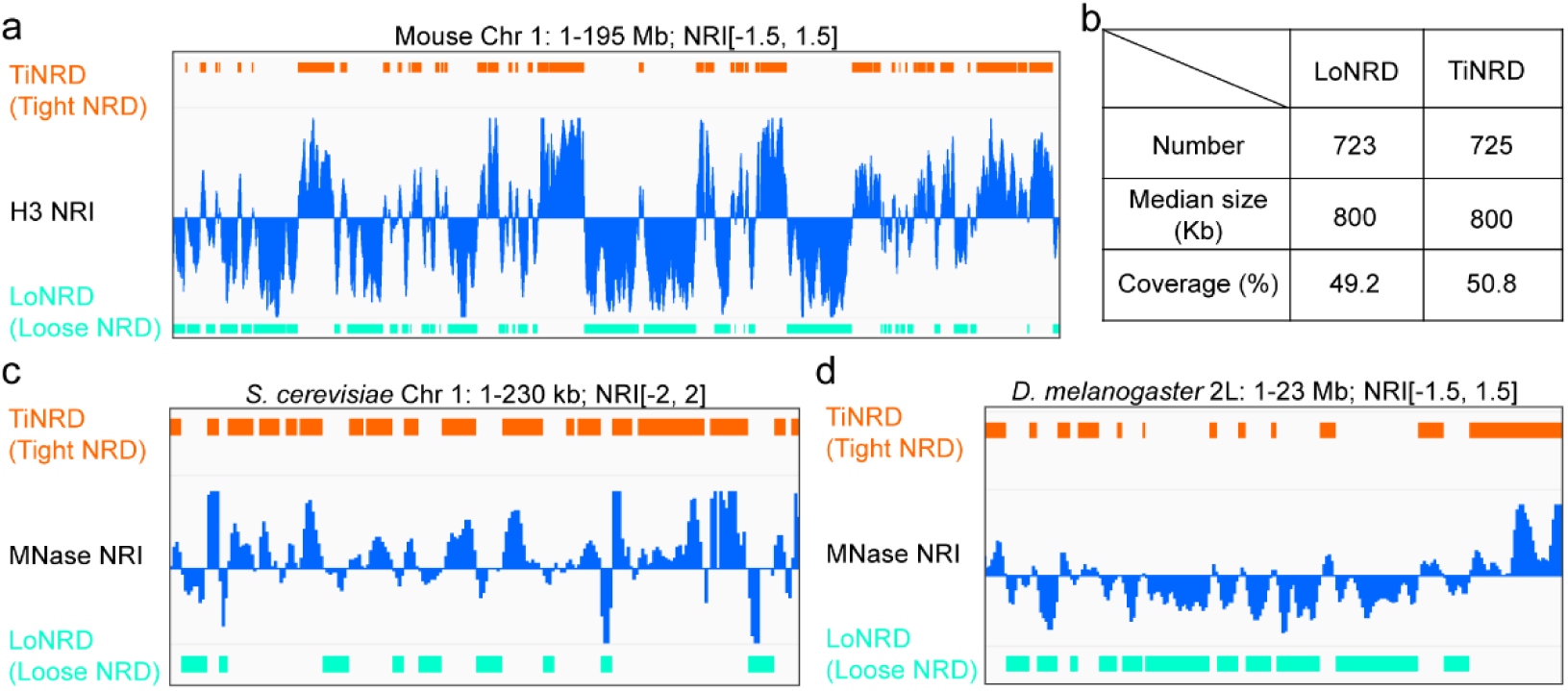
Nucleosome wrapping index is robust for nucleosome wrapping state characterization. a, Diagram shows the calculation of NRS, “a” is the number of fragments within X-250 bp and “b” is the number of fragments within 50-X bp. b-c, IGV tracks show NRS (b) or NRI (c) of chromosome 1 when different break point is chosen, including 80 bp, 90 bp, 100 bp, 110 bp, 120 bp, 130 bp, 140 bp, 150 bp, 160 bp. d, IGV tracks show the similarity between NRIs of H3 ChIP, H4 ChIP or MNase input on Chromosome 1. e, IGV tracks show the dynamic changes of NRSs and NRIs under time-course MNase digestion on Chromosome 1. f, Dot plot shows the strong positive correlation between H3 NRI and H3 genome coverage. “r” indicates the Pearson correlation coefficient. g, IGV tracks show NRIs, MNase input signals and genome coverage signals of sonicated genomic DNA from G1, S and G2/M phase sorted mouse ES cells on Chromosome 1.

When NRI(140) was calculated for histone H4 wrapping-seq data and the MNase digestion input data, they both show high correlation with H3 NRI(140) (Fig. 1d, Fig. S1g), suggesting that MNase-seq data can be used to analyze nucleosome wrapping states. We further calculated NRS(140) and NRI(140) of MNase input date with 10∼50 min time-course digestion. We found that the genome-wide NRS(140) generally shift from positive value to negative value (Fig. 1e, Fig. S1h), consistent with decreased proportion of long DNA fragments and increased proportion of short DNA fragments with increasing digestion time. However, the NRI(140) patterns from different digestion time points are highly similar (Fig. 1e, Fig. S1i), and highly correlated (Fig. S1j), suggesting that MNase-seq NRI(140) is robust for nucleosome wrapping detection, despite of a certain degree of digestion variation.

Surprisingly, we observed that NRI is highly positive correlated with nucleosome density represented by H3 or H4 ChIP signal or MNase input signal (Fig. 1d, Fig. 1f). As the genome copy number, and thus the nucleosome density, can be varied as a result of DNA replication, we performed MNase wrapping-seq after sorting cells according to cell cycle phases of G1, S, G2/M. In parallel, we sequenced libraries prepared from sonicated genomic DNA of the sorted cells, so as to test whether MNase digestion *per se* would introduce nucleosome density variation. We found that in all three cell cycle phases, the signal of sonication input and MNase-seq input is highly similar viewed from IGV (Fig. 1g) and highly correlated at genome-wide (Fig. S1k), thus excluding that the variation of nucleosome density is a result of MNase digestion. Moreover, we found that nucleosome density indeed shows variation along the genome during S-phase, but not during G1 and G2/M phase (Fig. 1g). However, the NRI pattern are stably consistent among G1, S or G2/M phases (Fig. 1g). Thus, these results argue that the NRI pattern is not an artificial consequence of nucleosome density variation along the genome.

Taken together, these results demonstrate that quantification of nucleosome wrapping states by Nucleosome Wrapping Index is highly robust. Notably, this method is also very straightforward when taking advantage of MNase input data.

### Nucleosome wrapping domain is detected in yeast, fly and mouse genome

Interestingly, we observed from the genome-wide NRI pattern that there are large genome domains continue with characteristic nucleosome wrapping states (Fig. 1c, 1d, 1e, 1g). To qualify the nucleosome wrapping states along the genome, we segmented the genome based on NRI(140) of histone H3 (Fig. 2a). In total, the genome is resolved into 1448 <u>N</u>ucleosome Wrapping <u>D</u>omains (NRDs) (Fig. 2b). NRDs enriched with positive NRI were termed as tight NRDs (TiNRDs, n= 723); NRDs enriched with negative NRI were termed as loose NRDs (LoNRDs, n=725) (Fig. 2a, 2b). Both TiNRDs and LoNRDs in mouse ES cells have a median length of 800 Kb, and cover 49.2% and 50.8% of mouse genome, respectively (Fig. 2c). These NRDs maintain characteristic nucleosome wrapping features across multiple NRI groups (Fig. S2a), bin resolution (Fig. S2b) and histone types (Fig. 2c), suggesting that the qualification of nucleosome wrapping states by NRDs (refers to NRDs of H3 NRI(140) unless stated) is robust.

**Figure 2.**
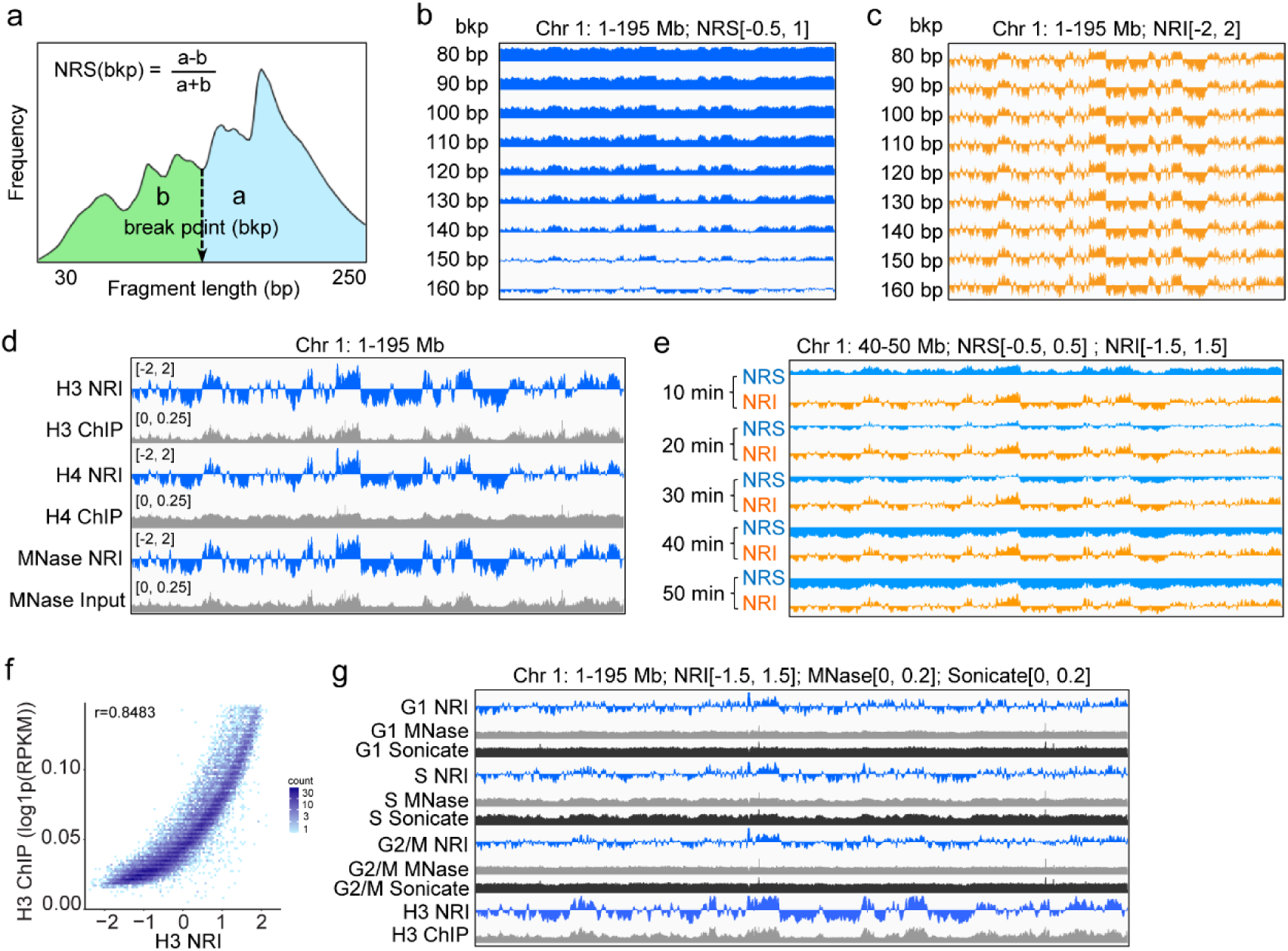
Nucleosome wrapping domain is detected in yeast, fly and mouse genome. a, IGV tracks show the distribution of TiNRDs (tight nucleosome wrapping domains) and LoNRDs (loose nucleosome wrapping domains) on Chromosome 1. b. Table shows the number, median length and genome coverage of TiNRDs and LoNRDs. c-d, IGV tracks show the distribution of TiNRDs and LoNRDs on Chromosome 1 of *S. cerevisiae* genome (c) or chromosome 2L of *D. melanogaster* genome (d).

To explore whether nucleosome wrapping domains is conserved in other species other than Mus musculus, we calculated NRI using MNase-seq data of *S. cerevisiae* ^16^ and *D. melanogaster* S2 cells ^17^, respectively. As in mouse ES cells, we observed stereotypical nucleosome wrapping domains in both S. cerevisiae (Fig. 2d, Fig. S2d) and *D. melanogaster* (Fig. 2e, Fig. S2e). Thus, these results suggest that regional nucleosome wrapping is conserved during evolution.

### Nucleosomes wrap tighter in euchromatin than in heterochromatin

To investigate the relationship between nucleosome wrapping and chromatin states, we computed the genome-wide correlation of H3 NRI(140) and DNase I sensitivity, H3K4me1, H3K4me3, H3K9me3, H3K27me3. We found that H3 NRI(140) is well positively correlated with DNase I sensitivity, and active chromatin markers such as H3K4me1 and H3K4me3, but not with heterochromatin markers H3K9me3 and H3K27me3 (Fig. 3a). We further counted the fragment length in the peak regions of H3K4me1, H3K4me3, H3K9me3 and H3K27me3. We found that euchromatin regions with H3K4me1 or H3K4me3 peaks show higher proportion of long DNA fragments than heterochromatin regions with H3K9me3 or H3K27me3 peaks (Fig. 3b), and that heterochromatin regions show higher proportion of short DNA fragments than euchromatin regions (Fig. 3b). Correspondingly, quantification of NRS in these peak regions showed that nucleosomes in H3K4me1 or H3K4me3 peaks regions have higher H3 NRS(140) that those in H3K9me3 or H3K27me3 peaks regions (Fig. S3a). On the other hand, when analyzing the epigenetic features of NRDs, we found that TiNRDs show higher level of euchromatin marks, such as H3K4me1, H3K4me3 than LoNRDs (Fig. S3b); while LoNRDs show higher level of heterochromatin marks such as H3K9me3 and H3K27me3 than TiNRDs (Fig. S3b). Together, these results suggested that nucleosomes wrap tighter in euchromatin than in heterochromatin.

**Figure 3.**
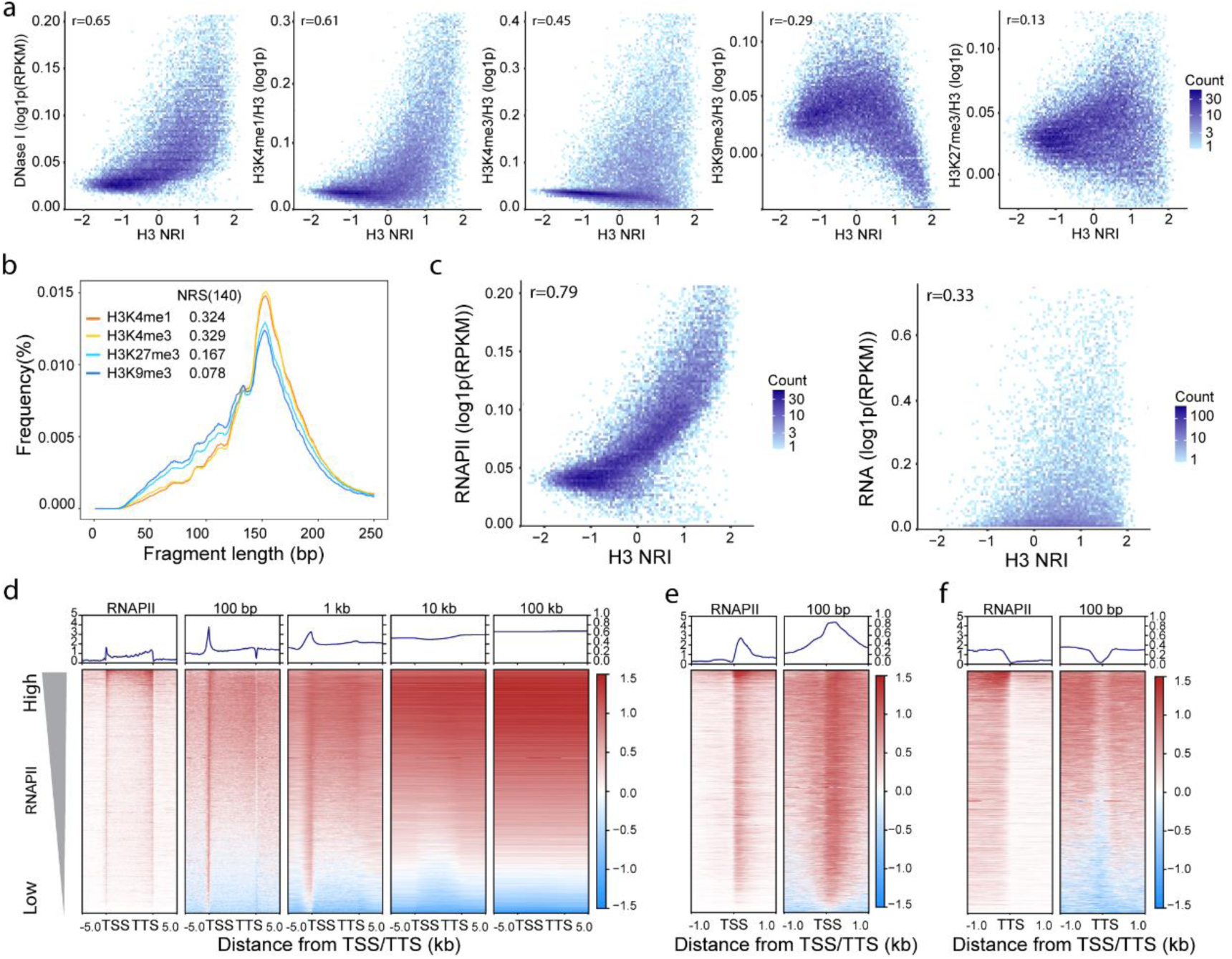
Nucleosomes wrap tighter in euchromatin than in heterochromatin. a, Dot plots show the genome-wide correlations between NRI (refers to NRI(140) of H3 unless stated) and DNase I, H3K4me1, H3K4me3, H3K9me3, H3K27me3. The signal of histone modifications was normalized by H3. b, Histogram shows the distribution of fragment length of histone H3 enriched DNA fragments in the peak regions of H3K4me1, H3K4me3, H3K9me3, H3K27me3, NRS(140)s is the nucleosome wrapping score calculated with the peaks of each markers. c, Dot plots show the correlations between NRI and Polr2a or total RNA-seq signal within gene body regions. d-f, Heatmaps show the distribution of H3 NRI at gene body regions (d), 2 kb regions around transcription start sites (TSSs) (e) and 2 kb regions around transcription termination sites (TTSs) (f). Genes were sorted descendently according to Polr2a signal in the gene body regions. H3 NRIs calculated with 100 bp, 1 kb, 10 kb and 100 kb bin resolution were shown for gene body regions. “r” indicates the Pearson correlation coefficient in a and c.

To explore the relationship between nucleosome wrapping and transcription activity, we computed the correlation between NRI and Polr2a ChIP signal or total RNA-seq signal at gene bodies. We found that NRI highly correlated with Polr2a ChIP signal, but to a weaker extend with total RNA level (Fig. 3c), suggesting that NRI is closely related to the RNA polymerase activity on the genes. After sorting the gene bodies according to Polr2a ChIP signal, we found that NRI decreased gradually with RNAPII signal (Fig. 3d, Fig. S3c). Moreover, NRI is highly enriched immediately downstream of TSS, and deceased within the gene bodies (Fig. 3d, 3e), suggesting +1 nucleosome is highly wrapped. This result is in consistent with that +1 nucleosome is involved in the regulation of RNA Polymerase initiation ^18,19^. At the transcription termination sites (TTSs), we found that NRI is low at the center of TTS, but it increased downstream beyond the TTS (Fig. 3f), suggesting there are other mechanisms regulating nucleosome wrapping beyond transcription elongation.

### Transcription promotes nascent nucleosome wrapping

During DNA replication, the nucleosome is dis-assembled ahead of the DNA replication fork, and re-assembled behind the replication fork. To study the nucleosome wrapping dynamics during DNA replication, we pulse-labeled mouse ESCs with EdU to label nascent nucleosomes assembled immediately behind the replication fork, then chased with thymidine to follow the maturation dynamics of nascent nucleosomes (Fig. 4a). Then we performed wrapping-seq using the bulk nucleosomes to calculate nucleosome wrapping states of mainly parental nucleosomes, and using the EdU labeled nucleosome to analyzed nucleosome wrapping states of the nascent nucleosomes. We found that the genome-wide NRI is highly correlated and stably between the parental nucleosomes and nascent nucleosomes at genome-wide (Fig. S4a) and in NRDs (Fig. S4b), suggesting that the NRDs are stably inherited during DNA replication. As in each “Nascent” or “Chase” condition, the “Input” (parental) nucleosomes and “Pulldown” (nascent) nucleosomes are from the same MNase digestion condition, the nucleosome wrapping score should be comparable directly. Interestingly, we observed that while the NRS of nascent nucleosomes is smaller than that of parental nucleosome under nascent condition, the NRS of the nascent nucleosomes became higher than that of parental nucleosomes under chase condition, in both TiNRDs and LoNRDs (Fig. 4b). This result suggested that, after DNA replication, the DNA wraps looser on the nascent nucleosomes than on the parental nucleosomes; however, within a time window, the nascent nucleosomes will mature and the DNA would wrap tighter than parental nucleosomes. This dynamic wrapping states of nascent nucleosomes can also be observed in the MINCE dataset of Drosophila S2 cells ^17^ (Fig. S4c). Together, these results suggested that the wrapping state of nascent nucleosomes is gradually matured after DNA replication, and this process is independent of the local nucleosome wrapping domains.

**Figure 4.**
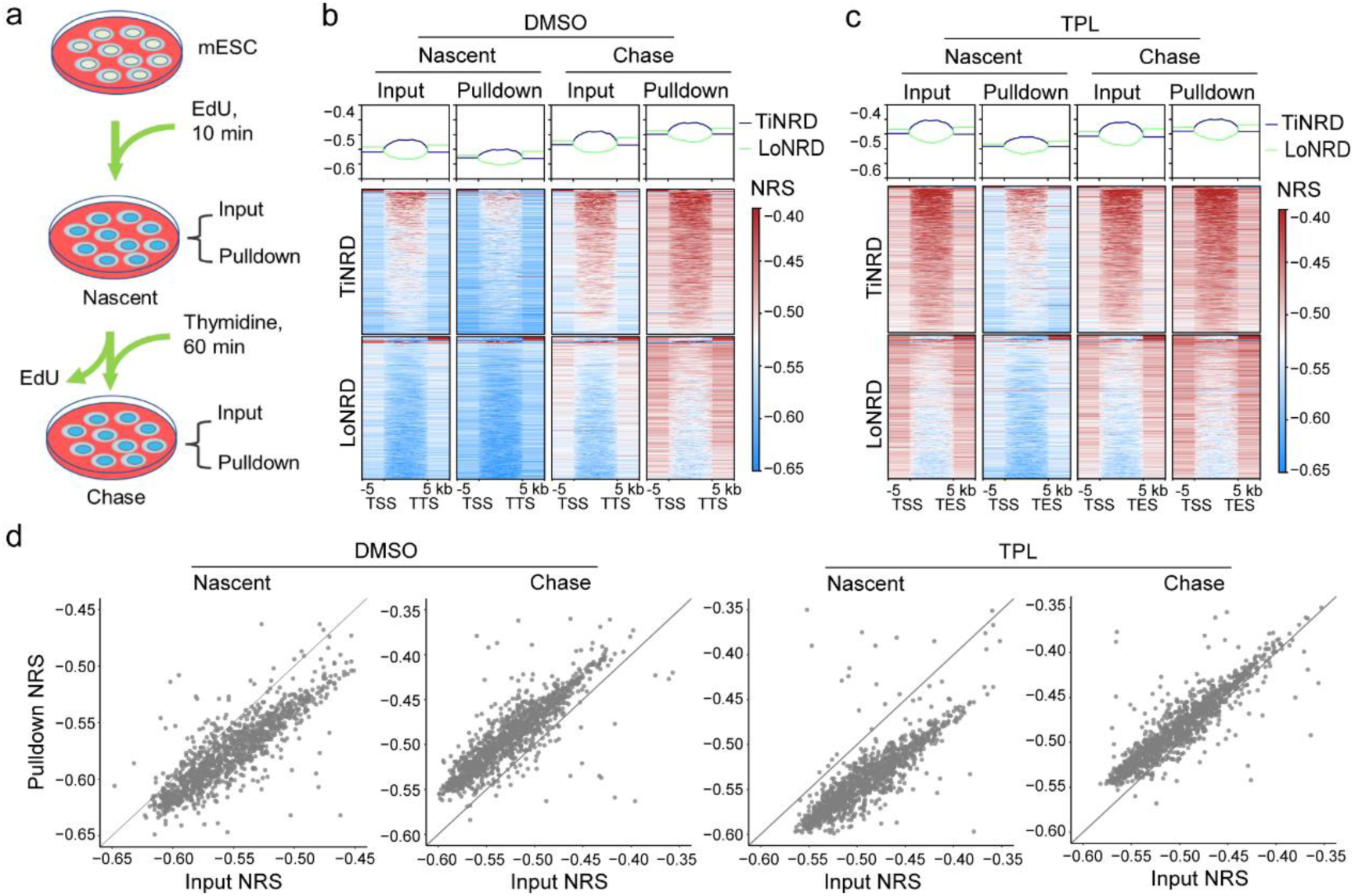
Transcription promotes nascent nucleosome wrapping. a, Diagram show the preparation of nascent and chase samples for nucleosome wrapping analysis during DNA replication. b-c, Heatmaps shows MNase NRS(140)s of parental and nascent nucleosomes from nascent and chase conditions, without (b) or with (c) triptolide (TPL) treatment. TPL was used to inhibit transcription initiation, and DMSO treatment was control. MNase NRS(140) was calculated using MNase-seq dataset with 100 kb bins and break point at 140 bp. d, Dot plots show the difference of NRS between parental and nascent nucleosomes as distance to the diagonal. TPL was used to inhibit transcription initiation, and DMSO treatment was control.

To explore the whether transcription regulates nucleosome wrapping, we inhibit transcription with triptolide (TPL) ^20^ during DNA replication. We found that, the establishment of NRDs are not affected by transcription inhibition (Fig. S4d). However, when analyzing NRS, we found that, under nascent condition, the NRS of nascent nucleosomes became more smaller than parental nucleosomes after TPL treatment (compare “Nascent” conditions in Fig.4b and Fig. 4c). Under chase condition, the NRS of nascent nucleosomes became less larger than parental nucleosomes after TPL treatment (compare “Pulldown” conditions in Fig.4b and Fig. 4c). Quantification the difference of NRS between nascent and parental nucleosomes as position relative to the diagonal in dot plot (Fig. 4d), or quantification of the difference of NRS by subtracting the NRS of parental nucleosome from the NRS of nascent nucleosome in NRDs as boxplot (Fig. S4e) confirmed these observations. Together, these results suggest that TPL inhibited the wrapping of nascent nucleosomes assembled behind the replication fork, and in other words, transcription promotes nascent nucleosome wrapping after DNA replication.

### Nucleosome wrapping states delineate Hi-C compartment domains and replication timing domains

It has been reported that the Hi-C compartments are highly resemble of replication timing domains ^21,22^. Surprisingly, we found that the genome-wide track of NRI is almost identical to the PC1 value ^23^ and the replication timing value of mouse ESC (Fig. 5a, Fig. S5), with TiNRDs correspond to A compartments and early RT domains, LoNRDs correspond to B compartments and late RT domains. Genome-wide correlation between NRI, PC1 and RT value also supports the observation (Fig. S6a), albeit with slightly higher correlation between H3 NRI and PC1 value than correlation between H3 NRI and replication timing value.

**Figure 5.**
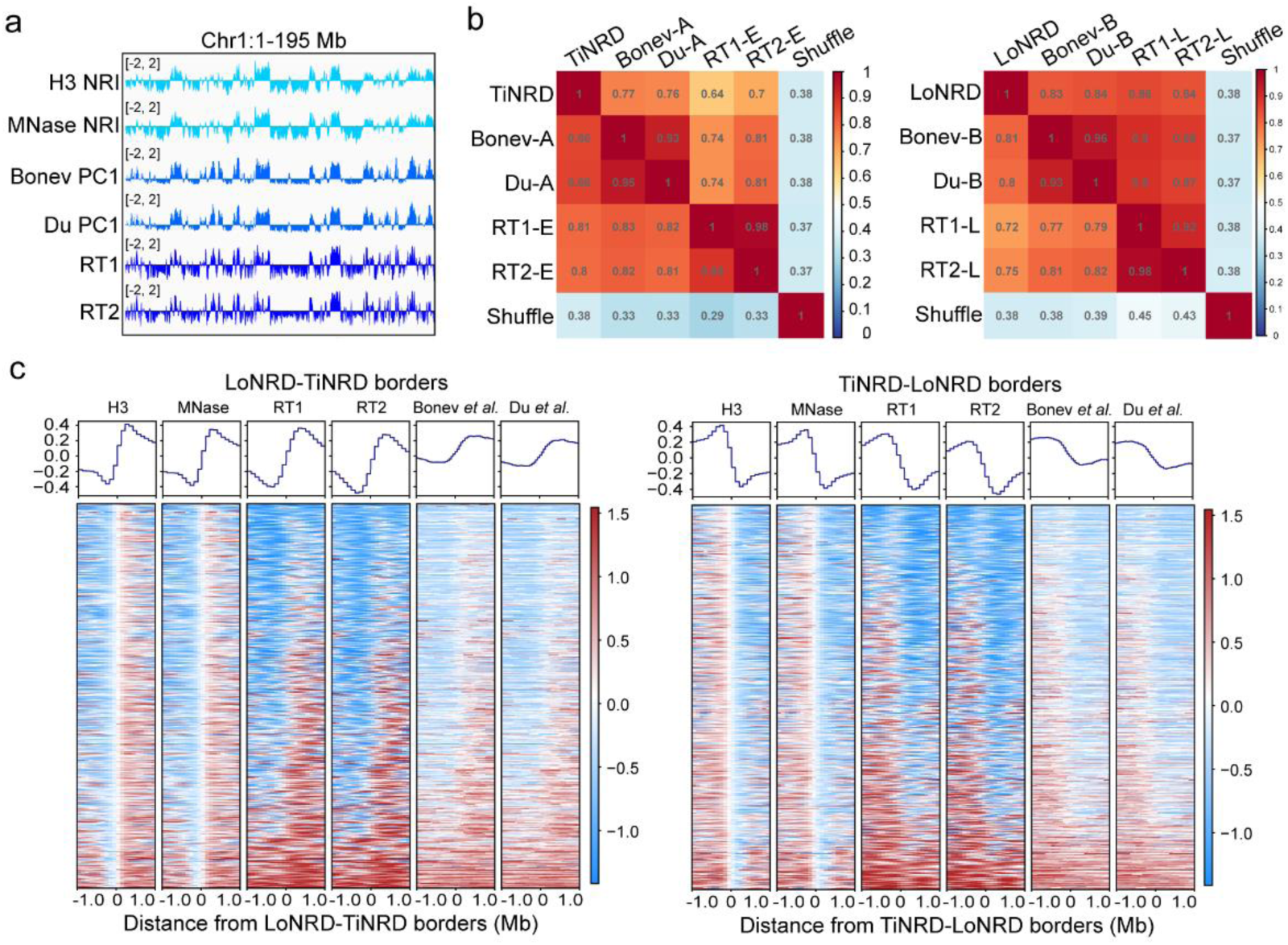
Nucleosome wrapping states delineate Hi-C compartment domains and replication timing domains. a, IGV tracks show the pattern of H3 NRI(140), Hi-C PC1 value and replication timing values on Chromosome 1. b, Heatmaps show the percentage of overlapping between NRDs and Hi-C compartment domains, RT domains. Values show the ratio of domains in the column direction to domains in the row direction. c, Heatmaps show the distribution of H3 NRI(140), MNase NRI(140), Hi-C PC1 values and replication timing values on the 2 Mb regions around LoNRD-TiNRD borders (left panel) or TiNRD-LoNRD borders (right panel).

We used NRDs with 100 kb resolution to calculated the percentage of region overlapping between NRDs, Hi-C compartments and replication timing domains. We found that 86% of the length of A compartments and about 80% of the length of early replication domains overlay with TiNRDs; about 80% of the length of B compartment and over 70% of the length of late replication domains overlay with LoNRDs (Fig. 5b). Moreover, at the LoNRD-TiNRD boarders, the Hi-C PC1 value and replication timing value sharply transit from negative to positive; reversely, at the TiNRD-LoNRD boarders, the Hi-C PC1 value and replication timing value sharply transit from positive to negative (Fig. 5c). Similar trend of NRI can be observed at the compartment boarders (Fig. S6b) and replication timing boarders (Fig. S6c). These results support the corresponding relationship between NRDs, Hi-C compartments and replication timing domains.

## Discussion

In this study, we have demonstrated that NRI can be utilized as a robust quantitative metric for characterizing the average wrapping states of nucleosomes within local chromatin regions. This quantification can be directly derived from MNase digestion sequencing data, independently of histone ChIP, and is not affected by the extent of MNase digestion. We further revealed the presence of non-uniform nucleosome wrapping states throughout the entire mouse genome, with a mutually spaced distribution of TiNRDs and LoNRDs. TiNRDs correspond to Hi-C A compartments and early replication timing domains, whereas LoNRDs correspond to B compartments and late replication timing domains. Thus, it’s interesting to deduced that, as depicted in the model (Fig. 6), in active chromatin regions represented by A compartments, nucleosome wrapping is relatively tight; whereas in repressive chromatin regions represented by B compartments, nucleosome wrapping is relatively loose.

**Figure 6.**
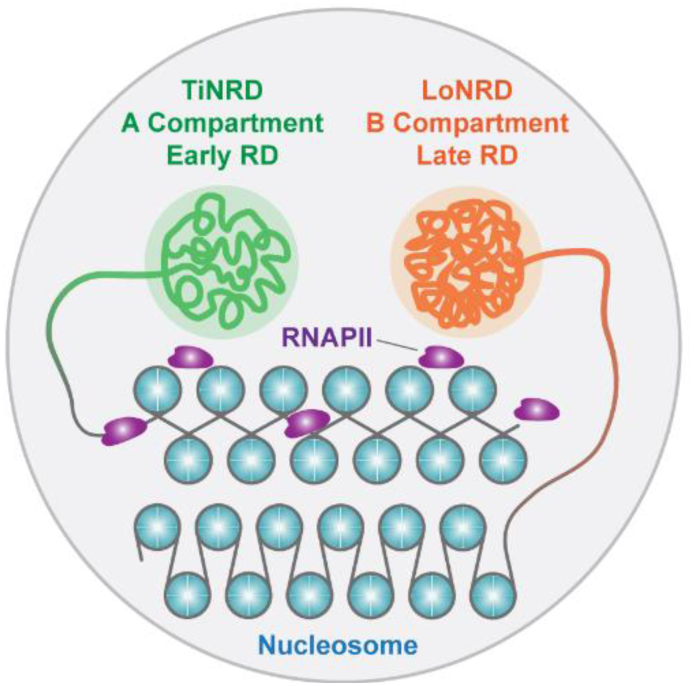
Nucleosome wrapping state encodes principles of 3D genome organization. Working model shows that genome of mouse ES cells is organized into nucleosome wrapping domains (NRDs), including TiNRDs and LoNRDs. TiNRDs and LoNRDs represents genomic regions enriched with relatively tight or loose wrapped nucleosomes, respectively. TiNRDs overlap precisely with Hi-C A compartment domains and early replication domains (RDs). LoNRDs overlap precisely with Hi-C B compartment domains and late replication domains. Thus, our model reveals a novel connection between genome organization at nucleosome array level and higher order level. Moreover, our model suggests that nucleosomes in active chromatin environment (A compartment) wrap tighter than nucleosomes in inactive chromatin environment, as a consequence of transcription promoted nucleosome wrapping.

Our data suggested that nucleosome wrapping becomes more relaxed after DNA replication and gradually tightens over time. Importantly, active gene transcription promotes the wrapping of nucleosome (Fig. 6). After transcription, nucleosomes need to be reassembled on the transcribed DNA to maintain chromatin structure. This reassembly is regulated by various histone chaperones and chromatin assembly factors ^24^. One of these histone chaperones that play central roles in nucleosome recycling during transcription is FACT (Facilitates Chromatin Transcription). FACT can bind histones and regulate the assembly or dis-assembly of nucleosomes to facilitate the passing of RNA Pol II on chromatin template ^25,26^. It’s recently reported that while the SPT16 subunit of the FACT complex can displace H2A-H2B dimers from the nucleosome and facilitate the formation of an open nucleosome structure, the SSRP1 subunit holds the H3/H4 tetramer on DNA and promoting the deposition of the H2A/H2B dimer onto the nucleosome, thus maintains the nucleosome integrity ^10,11,27^. These studies indicates that FACT may regulates the wrapping dynamics of nucleosome during active transcription.

While we have initially proved that transcription activity enhances the wrapping of nucleosomes in the genic (euchromatin) regions, the mechanisms regulating the nucleosome wrapping states in heterochromatin remains to be explored. The NuRD (Nucleosome Remodeling and Deacetylase) complex is required for specific gene silencing in heterochromatin regions. Through the ATP-dependent remodeling activity of the CHD subunit, it alters chromatin structure and DNA accessibility and regulates gene transcription ^28^. CHD3 is one the nine members of CHD family of ATPases (CHD1–9). It was recently reported that binding of CHD3’s tandem PHD fingers to histone H3 tails, which can be enhanced by H3K9me3 modification, promotes the unwrapping of nucleosomes *in vitro* ^29^. Heterochromatin protein 1 (HP1) is implicated in the formation and maintenance of heterochromatin via interaction with H3K9me3 modified nucleosomes ^30^. It was recently shown that binding of multiple Swi6 (the *S. pombe* HP1 protein) molecules to nucleosomes can promote HP1-chromatin LLPS (liquid–liquid phase separation), through increasing the solvent exposure of buried nucleosomal core histones ^31^. This HP1 mediated nucleosome reshaping process intrinsically resembles nucleosome unwrapping, indicating that HP1 binding may promote DNA unwrapping from the nucleosome core. While heterochromatin is characterized by its repressive nature, it is important to recognize that it is not a static and entirely silenced region of the genome. Instead, it is dynamically transited between repressive and active states, which involves various regulatory elements and factors. Thus, we speculate that the relative loose wrapping state of nucleosomes in the heterochromatin regions will allow controlled accessibility of regulatory factors to their target binding sites, during various cellular activities, such as gene expression regulation during development.

In summary, our research revealed a novel level of higher-order chromatin organization, which is intriguingly originates from the fundamental structural unit of chromatin—the nucleosome itself. This concept invokes the encoding of protein tertiary structure in amino acid sequences.

## Material and Methods

### Cell culture

Mouse ES cells (R1) were cultured in the medium with 80% DMEM (Thermo, 11965092), 15% FBS (Hyclone, SH30070.03), Nonessential amino acids (EmbryoMax, TMS-001-C), 2-Mercaptoethanol (EmbryoMax, ES-007-E), L-glutamine (EmbryoMax, TMS-002-C), Nucleosides (EmbryoMax, ES-008-D), Pen/Strep (EmbryoMax, TMS-AB-2C) and 1000U/ml leukemia inhibitory factor (LIF) (ESGRO, ESG1107) in standard incubator with 5% CO2 at 37 °C. For wrapping-seq, mouse ES cells plated for about 48 hours were directly crosslinked with 1% formaldehyde in PBS for 10 min at room temperature, then quenched by 125 mM glycine, and scraped off from the dish with a cell scraper. Then the cells were washed with cold PBS for twice and the cell pellet were stored at -80 °C.

To label nascent nucleosome, EdU (5-ethynyl-2’-deoxyuridine) was added at 20 μM final concentration and cultured for 10 min. Then the medium with EdU was drained off, and the cells were washed with 1XPBS pre-warmed at 37 °C for twice: to prepare nascent sample, the cells were crosslinked directly as described above for wrapping-seq; to prepare chase samples, the cells were further culture with mouse ES medium with 10 mM thymidine for 60 min, then crosslinked directly as described above for wrapping-seq. To inhibit transcription initiation, mouse ES cells were cultured with 1 mM triptolide (Sigma, T3652) final concentration or equal volume of DMSO only as control for 45 min before EdU labeling, then triptolide or DMSO was added back during chasing.

### Wrapping-seq

#### 1. Wet lab experiments

Cell pellet of ∼1×10^7 cells was resuspended in 1 mL nuclei extraction buffer (10 Mm Hepes, PH 7.5; 300 mM NaCl; 3 mM MgCl2; 0.5% IGEPAL CA-630; 0.1% SDS) with EDTA-free protease inhibitor cocktail (Roch, 05892791001) and incubate on ice for min. Then cell pellet was washed once with buffer A (10 mM Tris, PH 7.5; 10 KCl; 60 mM NaCl; 3 mM MgCl2) with EDTA-free protease inhibitor cocktail. For MNase (Sigma, N3755) digestion, cell pellet was then resuspended in buffer A with 2 mM CaCl2 and 0.4 U/mL MNase, and incubated at 37 °C on a metal incubator shaking at 900 rpm for 30 min or as indicated for time-course digestion. 10 mM EDTA and 1% SDS at final concentration was added to stop the digestion.

For MNase-seq, the digested chromatin was crosslink reversed by heating at 65 °C for 6 hours, and DNA were extracted using a standard phenol-chloroform extraction procedure. For paired-end sequencing, libraries without size selection were prepared as described in ^16^ using NEBNext Ultra DNA Library Prep Kit for Illumina (E7370L) and were sequenced on a DNBSEQ-T7 (MGI) platform with PE150 model.

For H3 or H4 ChIP, the digested chromatin was sonicated using a Q800R3 Sonicator (Qsonica), and the chromatin particles before crosslink reversion are about 1000-2000 bp after resolved on native 1% agarose gel, which is the typical chromatin particle size for ChIP-seq. Then, chromatin was first incubated with H3 (Abcam, ab1791) or H4 (Cell Signaling Technology, 14149S) antibody in RIPE-150 (50 mM Tris-HCl, 150 mM NaCl, 1 mM EDTA, 0.5% Triton X-100, protease inhibitors) at 4°C, then the BSA blocked protein A/G Dynabeads were added and incubated overnight at 4°C. The Dynabeads were washed by RIPE-150 with 0.1% SDS for 5 times, and eluted with Direct Elution Buffer (10mM Tris-HCl pH8, 0.3M NaCl, 5mM EDTA pH8, 0.5%SDS). The chromatin was crosslink reversed by heating at 65 °C for 6 hours, and DNA were extracted using a standard phenol-chloroform extraction procedure. For paired-end sequencing, libraries without size selection were prepared as described in ^16^ using NEBNext Ultra DNA Library Prep Kit for Illumina (E7370L) and were sequenced on a DNBSEQ-T7 (MGI) platform with PE150 model.

#### 2. Sequence data alignment

Paired-end reads were trimmed for adaptor sequence using cutadapt v4.3 ^32^ with parameters: -a AGATCGGAAGAGCACACGTCTGAACTCCAGTCAC -A AGATCGGAAGAGCGTCGTGTAGGGAAAGAGTGT -e 0.1 -n 2 -m 35 -q 30 –pairfilter = any, and then mapped to mm10 using Bowtie2 v2.5.1 ^33^ with parameters: -I 10 -X 1000 -3 5 -- local --no-mixed --no-discordant --no-unal. Duplicates were marked using picard MarkDuplicates v2.27.5 (https://broadinstitute.github.io/picard/) with default parameters and removed using samtools view v1.17 ^34^ with parameters: -f 2 -F 1024 -q 10. Unique pair-end reads in bam format was converted to bed format grouped according to fragment length using a custom python script.

#### 3. Nucleosome wrapping score (NRS) or Nucleosome wrapping index (NRI) calculation

Mouse genome was segmented in to non-overlapping 100 kb (or 100 bp, 1 kb, 10 kb) bins. Then, within each genomic bin, fragment from each fragment length group were counted by annotateBed from bedtools v2.30.0 ^35^. Fragment length groups used includes 50-80 bp, 80-90 bp, 90-100 bp, 100-110 bp, 110-120 bp, 120-130 bp, 130-140 bp, 140-150 bp, 150-160 bp, 160-250 bp or 50-140bp, 140-250bp. Then when a fragment length break point (X bp) is chosen, nucleosome wrapping score (NRS) is computed as the relative deviation between the number (a) of fragments within X∼250 bp and the number (b) of fragments within 50∼X bp, expressed as NRS(X) = (a-b)/(a+b). Then the genome-wide NRS was transformed as z-score to derive genome-wide nucleosome wrapping index (NRI). For IGV (Integrative Genomics Viewer) ^36^ visualization, NRS or NRI was Loess-smoothed by 10 bins using R function loess() from R package ‘stats’.

#### 4. Nucleosome wrapping domain (NRD) detection

FindHiCCompartments from homer2 ^37^ was used to detect tight NRD (TiNRD) based on the 10 bins smoothed 100 kb bin NRI (140bp) data of H3 with default parameter; and the parameter -opp was used to output the loose NRD (LoNRD). TiNRDs and LoNRDs were genomic domains with continuous positive or negative NRI, respectively. Genomic features with NRDs were annotated using annotatePeaks.pl from homer2 ^37^. Heatmaps were produced using plotHeatmap from deeptools v3.5.1 ^38^ for NRI or NRS visualization in NRDs or genic regions.

### Replication timing profiling for mouse ES cells

Replication timing profiling and timing domain for mouse ES cells was performed as described ^39,40^. Briefly, mouse ES cells were pulse labeled with 100 μM BrdU (5-Bromo-2’-deoxyuridine) for 2 hours, then trypsinized to single cell suspension and fixed with ice-cold ethanol. Then the cells were stained with Propidium iodide and cells in S-phase were FACS sorted into four fractions (S1, S2, S3, S4) according to Propidium iodide signal. Then genomic DNA was prepared for each sample via a standard phenol-chloroform extraction procedure, and then sonicated via a Q800R3 Sonicator (Qsonica) to average 200 bp fragment size. NGS adaptor were ligated using NEBNext Ultra DNA Library Prep Kit for Illumina (E7370L). Then the DNA were denatured by heating at 95 °C for 5 min and then cool on ice for 2 min. Nascent DNA strands were enriched via pulling down by anti-BrdU antibody (BD Pharmingen, cat. no. 555627). Library were amplified and sequenced on a DNBSEQ-T7 (MGI) platform with PE150 model. Sequence data alignment was performed as described above.

Sequencing data were aligned to mouse genome as described above. To calculate genome-wide replication timing, mouse genome was binned into non-overlapping 50 kb genome windows. Within each window, fragments from each sample were counted by annotateBed from bedtools v3.5.1 ^38^. After normalizing the total counts from each sample to 1 million, replication timing were calculated as log2((S1+S2)/(S3+S4))), and quantile normalized with R function normalize.quantiles.use.target() from R package ‘preprocessCore’. The quantile normalized replication timing value were used for replication timing domain detection via FindHiCCompartments from HOMER ^37^. Heatmap was produced using plotHeatmap from deeptools v3.5.1 ^38^. For IGV (Integrative Genomics Viewer) ^36^ visualization, the normalized value was Loess-smoothed using R function loess() from R package ‘stats’.

### Data availability

High throughput sequencing data generated in this study were deposited to Gene Expression Omnibus (GEO) under accession GSE243091.

Drosophila MINCE-seq fastq data was downloaded according to GEO record GSE76120, and mapped to drosophila genome version dmel-r6.16 (https://flybase.org). Samples GSM1974516 and GSM1974518 were merged as nascent input, and used for NRI visualization and NRD detection with 100 kb bin resolution as shown in Fig. 2d. Samples GSM1974517 and GSM1974519 were merged as nascent pulldown. Samples GSM1974528 and GSM1974530 were merged as chase input. Samples GSM1974529 and GSM1974531 were merged as chase pulldown. NRS was calculated for nascent input, nascent pulldown, chase input and chase pulldown samples as shown in Fig. S4c. *S. cerevisiae* MNase-seq fastq data was downloaded according to GEO record GSE30551, and mapped to yeast genome GCF_000146045.2 (https://www.ncbi.nlm.nih.gov). Sample SRR3193265 were used for NRI visualization and NRD detection with 1 kb bin resolution as shown in Fig. 2c.

ChIP-seq bam files and peak files of mouse ES cell line Bruce4, including H3K4me1, H3K4me3, H3K9ac, H3K9me3, H3K27ac, H3K27me3, Polr2a and total RNA-seq were downloaded under accession ENCSR343RKY from https://www.encodeproject.org. Hi-C compartments PC1 values of mouse ES cell was downloaded under accession 4DNESUCLJAZ8 ^41^ or 4DNESMXBLGKA ^23^ from https://data.4dnucleome.org.

All other data are available from the corresponding author upon reasonable request.

## Acknowledgments

This work was supported by National Key R&D Program of China (2022YFA1302801) and Shenzhen Science and Technology Program grant (RCYX20221008092930079) to HL. This work was also supported by the National Natural Science Foundation of China (32000423) and Shenzhen Science and Technology Program grant (GXWD20201231165807008, 20220817134430001) to ZW.

## Author contributions

Z.W. and H.L. designed and supervised the research; Z.W. performed experiments and bioinformatic analysis of wrapping-seq; X.Y. performed BrdU chasing experiments under supervision of Z.W.; R.F. performed replication timing experiments under the supervision of H.L.; Z.W. and H.L. prepared the manuscript with contributions from all authors.

---

**Figure S1.**
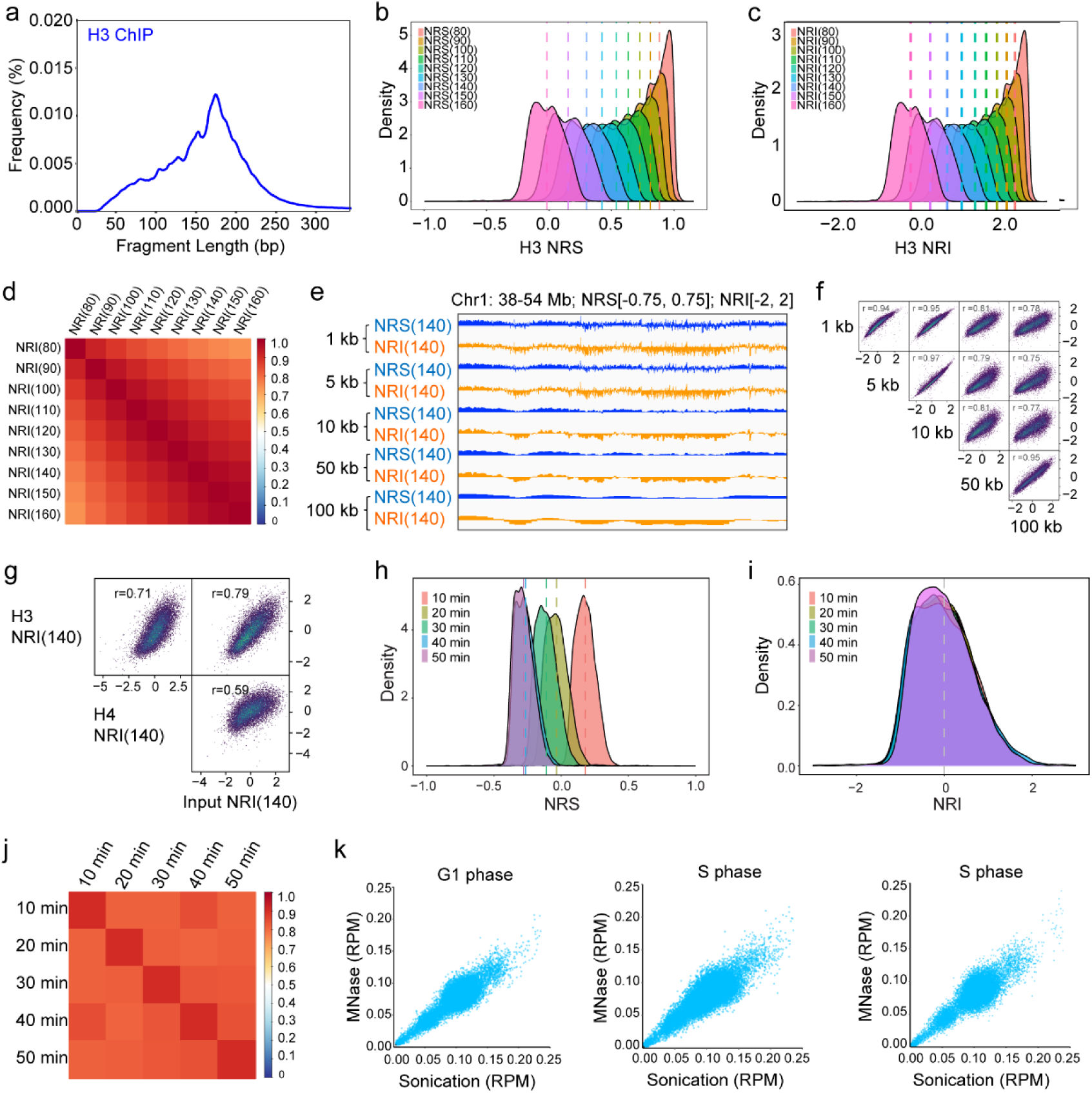
Nucleosome wrapping index is robust for nucleosome wrapping state characterization. a, Histogram shows the length distribution of DNA fragments enriched by histone H3 within 0-350 bp range. b-c, Histograms show the distribution of genome-wide NRSs (b) or NRIs (c) when different break point is chosen as in Fig. 1b. d, Heatmap shows the correlations between genome-wide NRIs when break point is chosen as in Fig. 1b. e, IGV tracks show NRSs and NRIs of H3 on Chromosome 1 calculated using different length of genome windows. f, Heatmaps show the genome-wide correlations between NRIs of H3 calculated using different length of genome windows. g, Heatmaps show the genome-wide correlations between NRI(140)s of H3, H4 or MNase input. “r” indicates the Pearson correlation coefficient. h-i, Histograms show the distribution of genome-wide NRSs (h) or NRIs (i) of MNase input under time-course MNase digestion. j, Heatmap shows the genome-wide correlation between NRIs of MNase input under time-course MNase digestion. k, Dot plots show the correlations between coverage of sonicated genomic DNA and coverage of MNase digested chromatin in G1, S and G2/M phases sorted cells.

**Figure S2.**
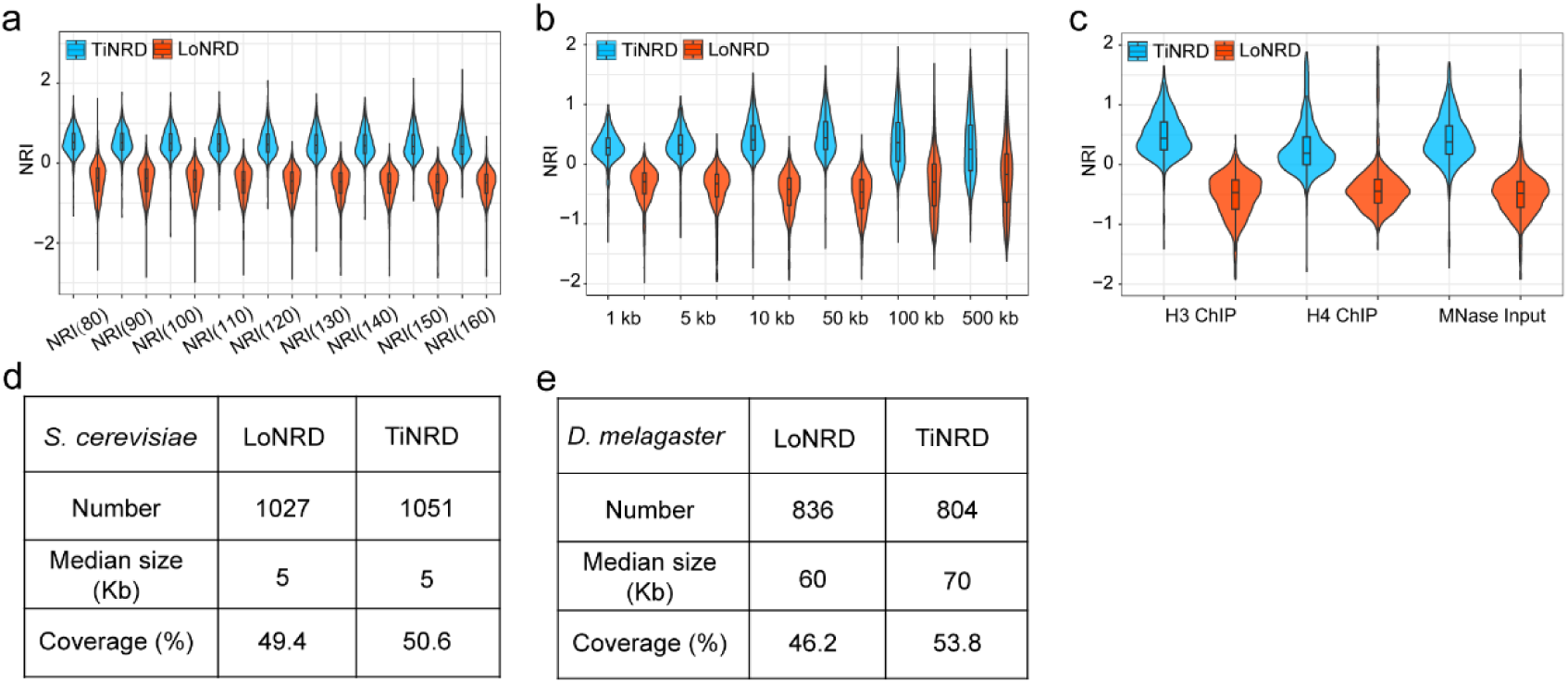
Nucleosome wrapping domain is detected in yeast, fly and mouse genome. a-c, Boxplots show the characteristic distribution of genome-wide NRIs in TiNRDs and LoNRDs. NRIs were calculated with different break points (a), bin lengths (b) or dataset (c). d-e, Tables show the number, median length and genome coverage of TiNRDs and LoNRDs of *S. cerevisiae* (d) and *D. melanogaster* genome (e).

**Figure S3.**
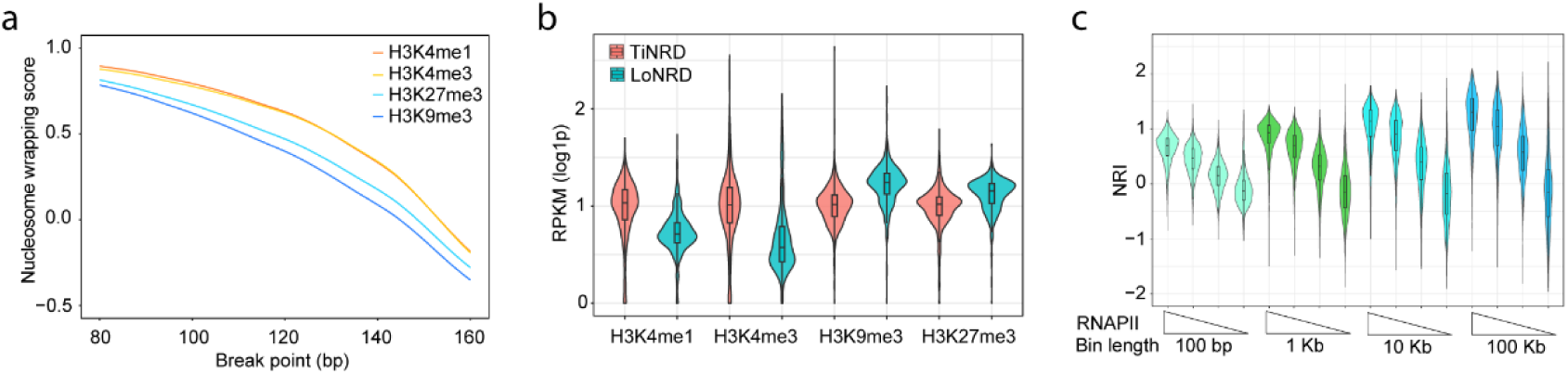
Nucleosomes wrap tighter in euchromatin than in heterochromatin. a, Line plot shows the H3 NRS(140)s of H3K4me1, H3K4me3, H3K9me3 and H3K27me3 calculated with H3 enriched DNA fragments in the corresponding peak regions, with break points range from 80-160 bp with 1 base pair step. b, Violin plots show the distribution of H3 normalized H3K4me1, H3K4me3, H3K9me3 and H3K27me3 signals in TiNRDs or LoNRDs. c, Violin plots show H3 NRIs in gene body regions grouped descendently by Polr2a signal. H3 NRIs calculated with 100 bp, 1 kb, 10 kb and 100 kb bin resolution.

**Figure S4.**
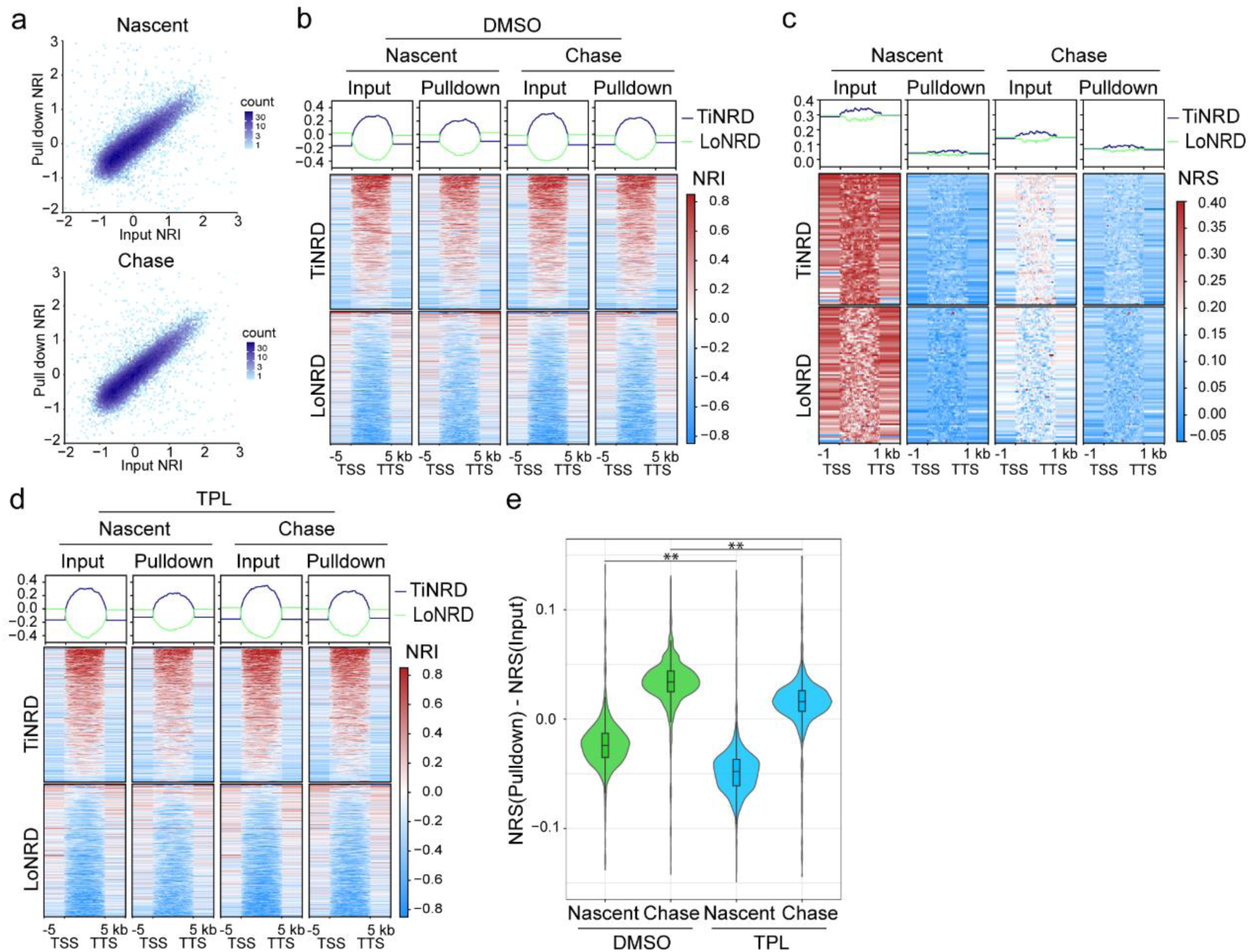
Transcription promotes nascent nucleosome wrapping. a, Dot plots show the genome-wide correlation between MNase NRI(140) of parental and nascent samples under nascent and chase conditions. MNase NRI(140) was calculated using MNase-seq dataset with 100 kb bins and break point at 140 bp. b, Heatmaps show MNase NRI(140) of parental and nascent nucleosomes from nascent and chase conditions. c, Heatmaps show MNase NRS(140) of parental and nascent nucleosomes from nascent and chase conditions, re-analyzed from MINCE-seq dataset {Ramachandran, 2016 #41}. d, Heatmaps show NRI of parental and nascent nucleosomes after TPL treatment. e, Violin plots show the dynamics of nucleosome wrapping after DNA replication, with or without TPL treatment. The difference of NRS was calculated by subtracting the MNase NRS(140) of parental nucleosome from the MNase NRS(140) of nascent nucleosome in each NRD. ** indicates P < 0.01.

**Figure S5.**
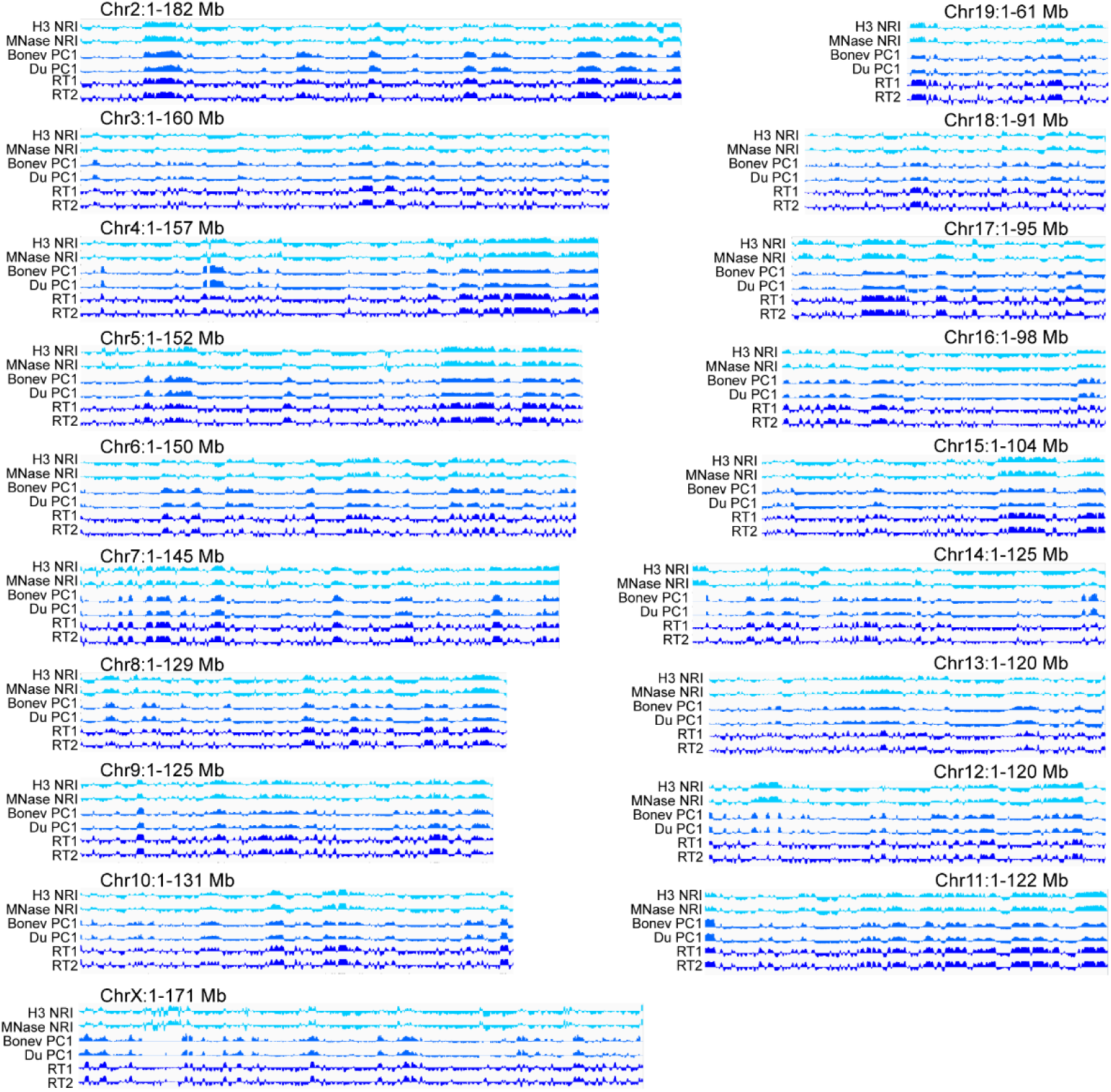
Nucleosome wrapping domains, Hi-C compartment domains and replication timing domains at mouse chromosomes. a, IGV tracks show per chromosome distribution of H3 NRI(140), MNase NRI(140), Hi-C PC1 values and replication timing values, except for chromosome Y.

**Figure S6.**
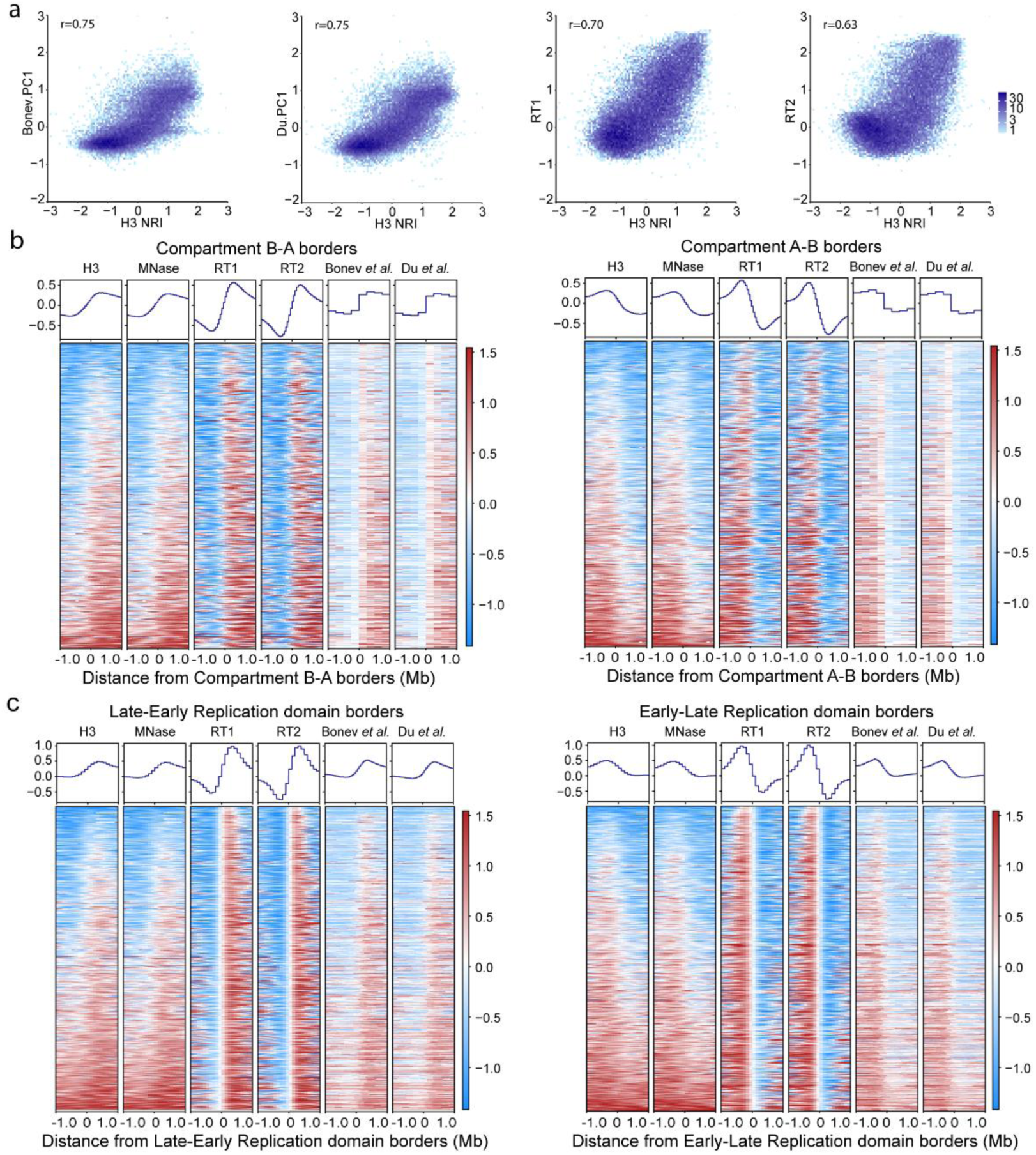
Nucleosome wrapping states delineate Hi-C compartment domains and replication timing domains. a, Dot plots show the genome-wide correlations between H3 NRI(140) and Hi-C PC1 values or replication timing values. “r” indicates the Pearson correlation coefficient. b-c, Heatmaps show the distribution of H3 NRI(140), MNase NRI(140), Hi-C PC1 values and replication timing values on the 2 Mb regions around B-A compartment domain borders (left panel) or A-B compartment domain borders (right panel) (b), or on the 2 Mb regions around late-early RT domain borders (left panel) or early-late RT domain borders (right panel) (c).

